# Signaling and transcriptional networks governing late synovial joint development

**DOI:** 10.1101/2021.08.25.457383

**Authors:** Qin Bian, Yu-Hao Cheng, Emily Y. Su, Yuqi Tan, Dong Won Kim, Hong Wang, Sooyeon Yoo, Seth Blackshaw, Patrick Cahan

**Affiliations:** Institute for Cell Engineering, Johns Hopkins School of Medicine, Baltimore MD 21205 USA; Department of Biomedical Engineering, Johns Hopkins School of Medicine, Baltimore MD 21205 USA; Department of Molecular Biology and Genetics, Johns Hopkins School of Medicine, Baltimore MD 21205 USA; Solomon H. Snyder Department of Neuroscience, Johns Hopkins School of Medicine, Baltimore MD 21205 USA

**Keywords:** single cell RNA-Seq, articular cartilage, ligament, synovium, chondrocyte, GRN

## Abstract

**Background:** During synovial joint development, cavitation marks the end of the emergence of new cell types and the onset of the consolidation of cell type specific programs. However, the transcriptional programs that regulate this crucial stage prior to joint maturation are incompletely characterized. Gdf5-lineage cells give rise to the majority of joint constituents such as articular cartilage, meniscus, ligament, and tendon. Therefore, to explore pre-maturation of the synovial joint, we performed single cell RNA-Seq analysis of 1,306 Gdf5-lineage cells from the murine knee joint at E17.5.

**Results:** Using computational analytics and *in situ* hybridization, we identified nine sub-states contributing to articular cartilage, meniscus, cruciate ligament, synovium, lining, and surrounding fibrous tissue. We identified a common progenitor population that is predicted to give rise to ligamentaocytes, articular chondrocytes, and lining cells. We further found that while a large number of signaling pathways orchestrate the differentiation of this progenitor to either ligamentocytes or to lining cells, only continued FGF signaling guides these cells to a default chondrocyte fate.

**Conclusions:** Our single cell transcriptional atlas is a resource that can be used to better understand and further study synovial joint development and the reactivation of embryonic programs in diseases such as osteoarthritis.

## 1. Introduction

Synovial joint degeneration is a major contributor to osteoarthritis, a disease with deep and broad impacts on human health in nations with increasing aging populations ^1,2^. New therapeutic approaches, such as cell replacement, and better models would benefit from an improved understanding of synovial joint development at different developmental stages ^3 4^. For example, we recently determined the transcriptional programs that regulate early (E12.5 to E15.5) synovial joint development by a combination of single cell RNA-sequencing (scRNA-Seq) of Gdf5-lineage joint progenitors, computational analyses, and in situ validation ^5^. While that study uncovered substantial transcriptional and fate bias heterogeneity in interzone cells, it did not characterize how joint progenitors ultimately commit to the major joint cell types, including permanent articular chondrocytes, ligamentocytes, and cells of the menisci and synovium. These lineages begin to be detectable around the time of cavitation, which in the hindlimb occurs around E16.5 ^6–8 9^.

To understand the transcriptomic programs active during late joint development, microarray have been applied to the E15-E16 elbow and knee joints ^10^, to the E16 meniscus ^11^. The transcriptomic characteristics identified by these investigations revealed the involvement of TGFβ (e.g. Gdf5) and Wnt (e.g. Sfrp2) signaling in knee morphogenesis, and the involvement of TGFβ (e.g. Lox) and IGF (Igf1) pathway in meniscus development. However, it is difficult to define the expression signatures of distinct sub-populations using bulk sample molecular profiling. The advent of single cell profiling makes it possible to achieve this aim ^12,13^. For example, a Lgr5^+^ population and a Tppp3^+^Pdgfra^+^ population were recently found that serve as progenitors for cruciate ligaments, synovial membrane, and articular chondrocytes ^14^, and for tendon ^15^, respectively.

To gain a comprehensive understanding of the transcriptional programs during late synovial joint development, we applied scRNA-Seq to Gdf5-lineage cells of the murine knee joint at E17.5. We combined computational analytics and *in situ* hybridization to identify the cell populations that contribute to different joint constituents and to uncover the transcriptional programs that mediate the lineage transitions. We found that Gdf5-lineage enriched cells consist of at least nine sub-populations that contribute to articular cartilage, meniscus, superficial lining, tendon/ligament, synovial fibroblasts, and other connective tissues. Furthermore, we predicted the signaling pathways, and the transcriptional programs underpinning the differentiation of joint progenitors to articular chondrocytes and to lining cells. We have made our data and analysis results available for the community to explore at https://e17-mouse.herokuapp.com

## 2. Materials and Methods

### Mice

We cross mated *Gdf5-cre* (Sperm Cryorecovery via IVF, FVB/NJ background, Jackson laboratory) mouse strain with B6.129×1-Gt(ROSA)26Sortm1(EYFP)Cos/J (RosaEYFP, gifted by the lab of Prof. Xu Cao from Johns Hopkins University) strain to generate Gdf5-cre::Rosa-EYFP mice. The genotype of the mice was determined by PCR analyses of genomic DNA isolated from mouse tails using the following primers: Gdf5-directed cre forward, 5’GCCTGCATTACCGGTCGATGCAACGA3’, and reverse, 5’GTGGCAGATGGCGCGGCAACACCATT3’ (protocol provided by Prof. David Kingsley, HHMI and Stanford University). All the protocols were approved by the institutional review board of Johns Hopkins University.

### Mice gender identification

We identified mouse gender by genotyping Sry Y gene using the primers: forward, 5’CTGGAAATCTACTGTGGTCTG3’, and reverse, 5’ACCAAGACCAGAGTTTCCAG3’.

### Cell isolation

Mice were kept in light-reversed room (light turns on at 10 pm and turns off at 10 am). Timing was determined by putting one male mouse and two female mice in the same cage at 9 am and separating them at 4 pm on the same day. We count that midnight as E0.5 stage. On E17.5, the pregnant mice were sacrificed by CO_2_ at 3 am. The cells were isolated using the protocol (Primary culture and phenotyping of murine Chondrocytes) with modification: The embryos were rinsed three times in PBS on ice. Two presumptive knee joints were isolated by transfemoral and transtibial division in a single 3 cm dish and incubated in digestion solution I (3 mg / mL collagenase D, DMEM high glucose culture medium, serum free) for 45 min at 37 °C, and then in digestion solution II (1 mg / mL collagenase D, DMEM high glucose culture medium, serum free) for 3 hrs (one embryo per dish) at 37 °C. During the period of incubation, the mice gender was identified by genotyping and only male samples were chosen for further processing. The tissues with medium were gently pipetted to disperse cell aggregates and filtered through 40 µm cell strainer, then centrifuged for 10 min at 400 g. The pellet was suspended with 0.4% BSA in PBS.

### Cell fractionation

All cells were fractionated by fluorescence-activated cell sorting (FACS). A MoFlo XDP sorter (Beckman Coulter, Miami, FL. USA) was used to collect YFP^+^ cells, and Propidium Iodide was used to exclude dead cells.

### Single cell RNA sequencing

GemCode™ Single Cell platform (10X Genomics) was used to determine the transcriptomes of single cells ^16^. Cells at 1000 / µl were obtained after sorting and placed on ice. One sample was selected and profiled based on the viability and amount. A total of 6000 cells were loaded, followed by GEM-RT reaction, and cDNA amplification. Single cell libraries were constructed by attaching P7 and P5 primer sites and sample index to the cDNA. Single cell RNA sequencing was performed on Illumina NextSeq 500 and HiSeq 2500 to a median depth of 168,000 reads per cell.

### Analysis and visualization of scRNA seq data

CellRanger (version 2.0.0) was used to perform the original processing of single cell sequencing reads, aligning them to the mm10 reference genome. We used the command line interface of Velocyto, version 1.7.3, to count reads and attribute them as spliced, un-spliced, or ambiguous ^17^. The resulting loom file was then subjected to quality control processing, normalization, estimation of cell cycle phase, clustering, and differential gene expression analysis using Scanpy 1.5.1 ^18^. Specifically, we excluded cells in which mitochondrial gene content exceeded 5% of the total reads or cells in with fewer than 501 unique genes detected. Then, we excluded genes that were detected in fewer than 3 cells, as well as mitochondrially-encoded genes, genes encoding ribosomal components, and the highly expressed lncRNA Malat1, resulting in a data set of 2,267 cells and 15,071 genes. Then, we performed an initial normalization on a per cell basis followed by log transformation, and scaling. We scored the phases of cell cycle using cell cycle-associated genes as previously described ^19^. Then we identified the genes that were most variably expressed across the whole data set, resulting in 2,176 genes. We performed PCA and inspected the variance ratio plots to determine the ‘elbow’, or number of PCs that account for most of the total variation in the data. We generated a graph of cell neighbors using diffusion maps ^20^, and then we performed Leiden clustering ^21^, which we visualized with a UMAP embedding ^22^. We also analyzed this with SingleCellNet ^23^, which had been trained using the Tabula Muris data set ^24^. Using a combination of SingleCellNet classification and manual annotation, we identified and removed non-joint cells as described in main text. Differentially expressed genes were identified using the Scanpy rank_genes_groups function. Gene set enrichment analysis was performed using GSEAPY (https://github.com/zqfang/GSEApy), a Python interface to enrichR ^25,26^. scVelo was used to compute cell velocities as previously described ^27^. CellRank was used to infer the starting and end states, and to compute the probability of each cell transitioning to each end state. We performed GRN analysis with Epoch ^28^ for each trajectory (i.e. progenitor 8 to chondrocyte, and progenitor 8 to lining cells) separately by first isolating those cells in progenitor cluster 8 and progeny clusters based on CellRank probability of reaching the selected terminal state, Cells within a trajectory were then ordered based on velocity pseudotime.

### Histochemistry and immunohistochemistry

The specimens were fixed in 10% buffered formalin for 24 hrs at RT, washed with distilled water and equilibrated in 30% sucrose in PBS at 4°C overnight, then mounted in O.C.T and frozen at -80°C. Ten-micrometer-thick coronal-oriented or sagittal-oriented sections were performed by cryostat. We performed Trichrome staining according to Trichrome Stain (Connective Tissue Stain) Kit protocol. Immunostaining was performed using a standard protocol. Sections were incubated with primary antibodies to mouse GFP (1:200) in Antibody Diluent, at 4 °C overnight followed with three 5 min washes in TBST. The slides were then incubated with secondary antibodies conjugated with fluorescence at room temperature for 1 h while avoiding light followed with three 5 min washes in TBST and nuclear stained with mounting medium containing DAPI. Images were captured by Nikon EcLipse Ti-S, DS-U3 and DS-Qi2.

### *In situ* hybridization

See KRT table for the information of oligonucleotides used for templates for antisense RNA probes. The T7 and SP6 primer sequence were added to 5- and 3-prime end, respectively. SP6 RNA polymerase was used for probe transcription. Probes were synthesized with digoxygenin-labeled UTP and hybridized at 68 °C overnight. Results were visualized by Alkaline phosphatase-conjugated anti-digoxygenin antibody and BCIP/NBT substrates.

### RNAscope Hiplex

RNAscope Hiplex probes were designed by Advanced Cell Diagnostics (ACD), Inc. Assay were performed according to ACD provided protocol as described in our previous study ^5^. See KRT table for details.

## 3. Results and discussion

### 3.1 Gdf5Cre+ cells in the knee joint from E17.5 Gdf5^Cre^R26^EYFP^ mouse are located in the superficial cartilage, ligament, menisci, synovium and non-joint tissues

To further study the heterogeneity of Gdf5-lineage cells at later stage of synovial joint development, we isolated YFP^+^ cells from the knee joint region of Gdf5Cre::R26EYFP (Gdf5EYFP) mice at E17.5 by enzymatic disassociation and fluorescence activated cell sorting (FACs) ^5^(**Supp Fig 1A,B**). A total of 2,648 cells were captured by the 10x Genomics Chromium platform and sequenced at a depth of 168,241 reads per cell (**Supp Table 1**). There were 2,267 cells remaining after removing potential doublets and low-quality libraries. We found nine clusters by the Leiden graph-based community detection algorithm ^21^ (**Supp Fig 2A**). We used a combination of automated cell-typing and marker gene expression to assign putative identity to the clusters. We removed cells representing types that do not contribute to major joint components, including hematopoietic cells (clusters 2, 3, 6, and 8), myoblasts (cluster 4), neural crest derived cells (Sox10 positive cells), endothelial cells and smooth muscle cells (**Supp Fig 2B-C**). After this process, a total of 1,306 Gdf5-lineage enriched (GLE) cells remained and were analyzed in depth. To localize these GLE cells, we applied IHC on knee joint sections of E17.5 Gdf5^Cre^R26^EYFP^ mice. YFP+ cells were detected in the superficial cartilage, cruciate ligament, menisci, synovium, and in the surrounding soft tissue, and a small number of cells were detected in the deeper zone of cartilage (**Fig 1A-B**).

**Figure 1:**
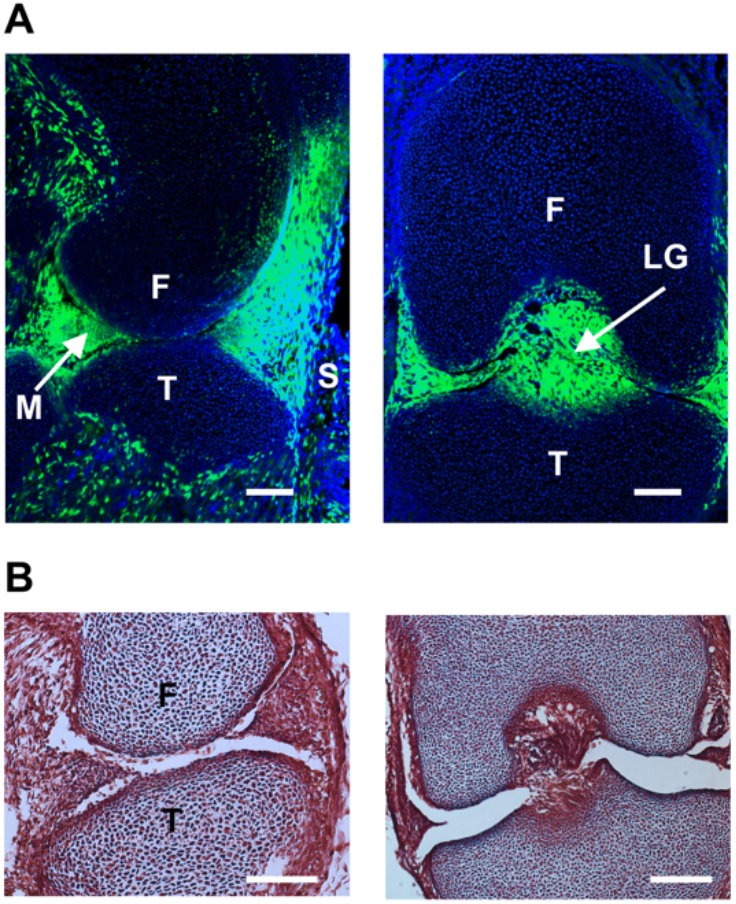
Localization of Gdf5-lineage cells. (A) IHC staining for GFP in sagittal section (left) and coronal section (right) of E17.5 knee joint. DAPI stains nucleus blue. (B) Morphology of E17.5 knee joint as indicated by Trichrome staining. Scale bar = 100 μM. F: Femur, T: Tibia, M: Meniscus, S: Synovium, CL: Crucial ligament

### 3.2 Transcriptional signatures define nine groups of GLE cells

We re-clustered the single cell data to determine the major transcriptional states and identities of the GLE cells. Using Leiden clustering, we found nine groups of cells (**Fig 2A**). To annotate these clusters, we used a combination of differential gene expression analysis and gene set enrichment analysis, followed by validation with ISH and RNAscope, as described below. Examining the genes preferentially expressed in each cluster immediately gave some hints as to their identity (**Fig 2B**). For example, we identified cluster 1 as chondrocytes by the expression of Col2a1, Col9a1, Col9a3, and the enrichment of the GO category “Cartilage development” (**Fig 2C**). Similarly, we identified cluster 2 as ligamentocytes based on the expression of Scx, Tnmd, and Thbs4 ^29^ and the enrichment of the GO category “Collagen fibril organization”, a prediction that we confirmed by RNAscope (**Fig 2D**).

**Figure 2:**
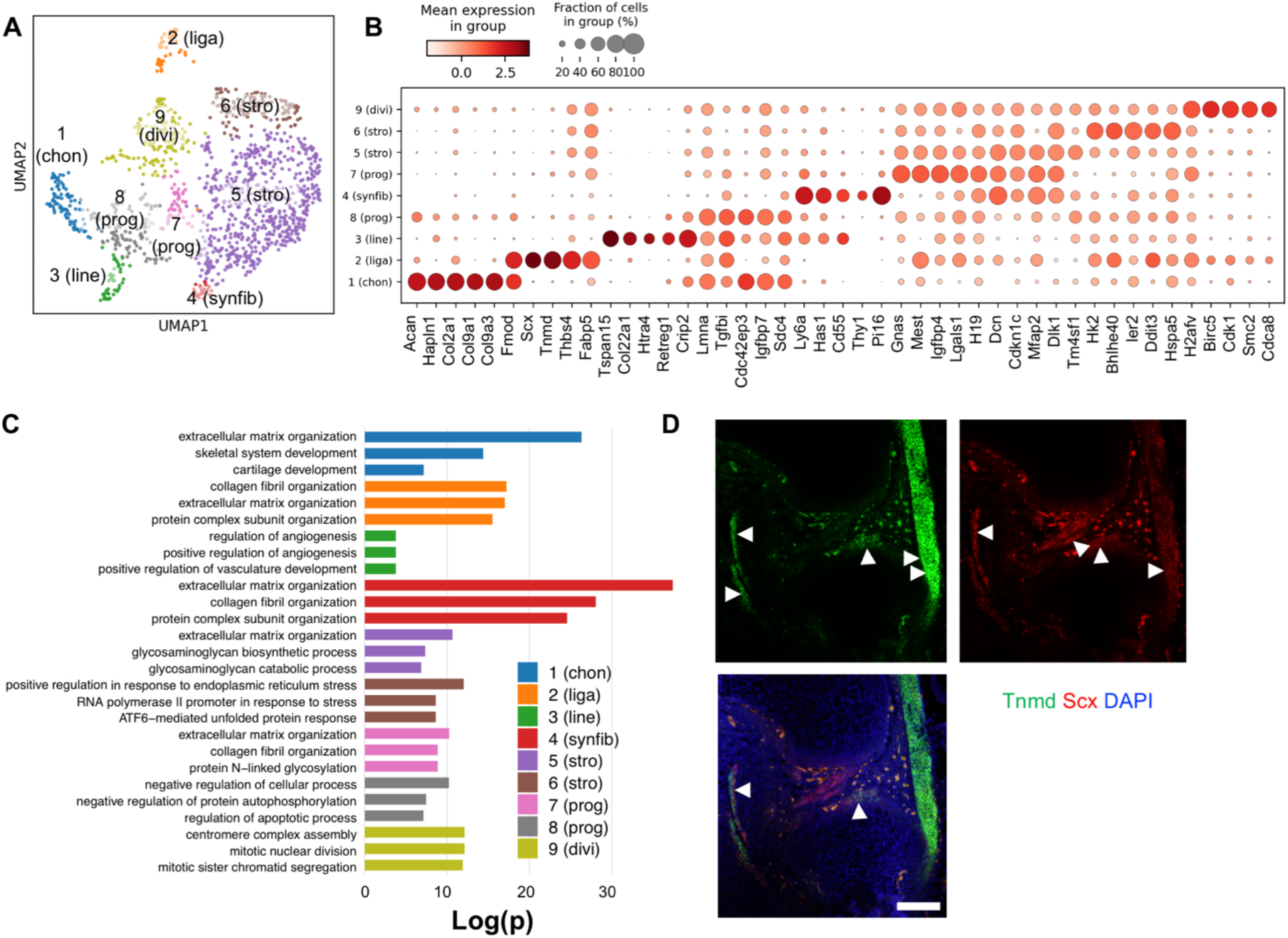
Identification of nine groups of GLE cells. (A) Leiden clustering and UMAP embedding of GLE cells, colored by cluster. (B) Dot plot of the top 5 differentially expressed genes in each cluster. (C) Top three enriched categories per group by gene set enrichment analysis. (D) Coronal sections of E17.5 knee joint showing expression of Tnmd (green) and Scx (red) marking cluster 2 (liga) cells. DAPI stains nucleus blue.

Cluster 3 cells uniquely expressed Col22a1 and Tspan15 (**Fig 3A**). Collagen XXII expression has been reported to be restricted to tissue junctions of muscle and articular cartilage ^30^. Similarly, an examination of Col22a1 at e16.5 also found that it was expressed at the superficial layer ^14^, suggesting that cluster 3 represented a population of superficial lining cells. To test this hypothesis, we performed ISH and RNAscope to determine where cluster 3 genes were expressed in the joint. We detected Col22a1 in a very thin fibrous sheath lining the cartilage surface and menisci by ISH (**Fig. 3B**). Tspan15, a gene encoding a member of the tetraspanin family of cell surface proteins, has very similar expression pattern as Col22a1. We found that Tspan15 was also expressed at the superficial layer of articular cartilage and meniscus (**Fig. 3B**). Taken together with the observation that cluster 3 also expressed Prg4 and Crip2 ^31^ (**Supp Fig 3A**), we conclude that the cells in this cluster comprise the lining or ‘skin’ of articular cartilage.

**Figure 3:**
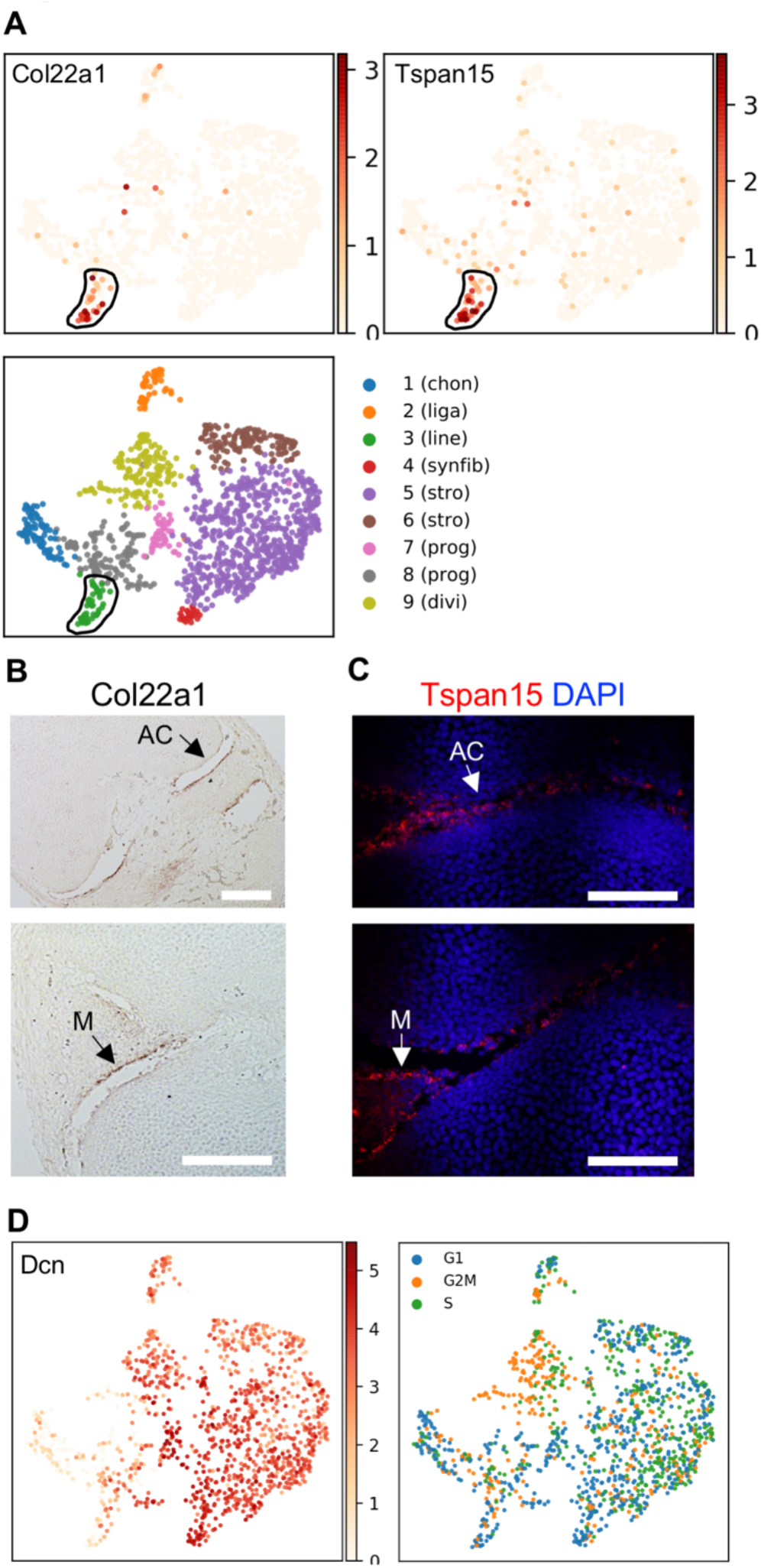
Cluster 3 is identified as a population of superficial lining cells. (A) Distribution of two Cluster 3 representative genes: Col22a1 and Tspan15. (B) ISH staining for Col22a1 and (C) RNAscope detection for Tspan15 of E17.5 knee joint sections. Arrows indicate positive staining at superficial layer of articular cartilage (upper panel) and meniscus (lower panel), Scale bar = 100 μM. AC: articular cartilage, M: meniscus. (D) Gene expression pattern of Dcn (left) and predicted phase of cell cycle in each group (right).

The preferential expression of fibroblast genes Dcn, Mfap2 ^32^, and Dlk1 in clusters 4, 5, 6, and 7 (**Fig 2B, Fig 3D**) supports the notion that these clusters are mesenchymal cells of the joint. Cluster 8 was made up of a mixture of proliferating chondrocytes and mesenchymal cells (**Fig 3D**).

Cluster 4 was marked by high levels of expression of Cd55, Thy1, and Has1 indicative of synovial fibroblasts ^33^. We note that many of these cells co-express genes that have previously be reported to distinguish inner synovial fibroblasts (Thy1) from synovial lining fibroblasts (Cd55, Has1). This discrepancy can be explained by species specific differences or in developmental timing differences between our data and prior reports.

The fact that no genes were substantially preferentially expressed in cluster 8 made it more challenging to identify. We noticed that it had detectable levels of genes that are preferentially expressed in chondrocytes (e.g. Acan and Cd9) (**Supp Fig 3B**), superficial lining cells (e.g. Crip1 and Crip2) (**Supp Fig 3C**), and in mesenchymal cells (e.g. Arl6ip5, Lmna, and Ptn) (**Supp Fig 3D,E**). This suggested that this cluster may represent a less-differentiated progenitor population expressing features of multiple downstream progeny. To determine the localization of this population, we examined the expression, *in situ*, of Col2a1, Acan, and Ptn, which collectively distinguished chondrocytes, stromal cells, and cluster 8 cells in our single cell data (**Fig 4A**). While Col2a1 was strongly expressed by chondrocytes in articular cartilage, we found that its expression was sparse and weak in menisci (**Fig 4B**). Although Acan expression was co-localized with Col2a1, its expression was evenly distributed in the inner menisci at a relatively low level as compared with its expression in the long bones. Ptn, on the other hand, was strongly expressed in menisci. Taken together, these observations support the notion that cluster 8 cells are found predominately in the meniscus at e17.5.

**Figure 4:**
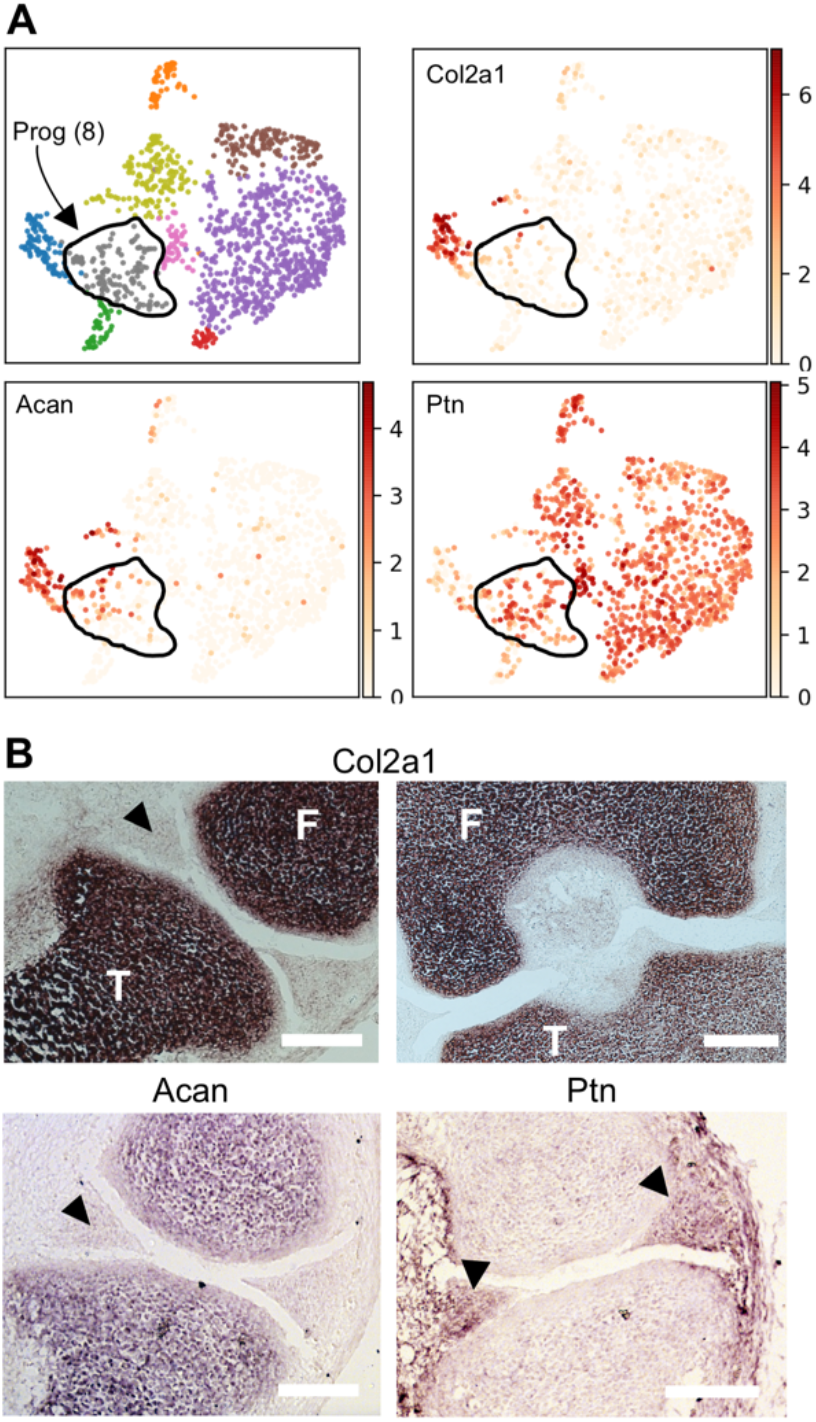
Cluster 8 is composed of fibro-chondrogenic progenitors that are localized at e17.5 in the meniscus. (A) Gene expression patterns of Col2a1, Acan, and Ptn. (B) IHC detection for Col2a1, Acan, and Ptn. Arrows point meniscus. Purple represents positive. Scale bar = 100 μM.

### 3.3 RNA velocity predicts common transcriptional origin of synovial fibroblasts, ligamentocytes, articular chondrocytes, and lining cells

Next, we performed RNA Velocity analysis to determine how the GLE cells were related to each other. In brief, RNA Velocity uses the ratio of spliced to un-spliced transcript counts to model transcriptional kinetics, which are then used to predict the future transcriptional state of each cell ^17,34^. Our application of RNA Velocity to e17.5 GLE cells revealed several general patterns of cell dynamics. First, we found that cells within each cluster were still undergoing dynamic transcriptional re-modeling (**Fig 5A**). Second, in most clusters, the velocities were unidirectional, for example, the chondrocyte velocities were pointing in the direction of higher Col2a1 and Col9a1 expression. Some clusters had unidirectional velocities towards another cluster, for example, the stromal cluster 5 largely had velocities ^35^ towards stromal cluster 6. Third, stromal clusters 7 and 8 clearly had velocities towards two or more other clusters, indicating that these clusters represented multi-potent progenitor populations.

**Figure 5:**
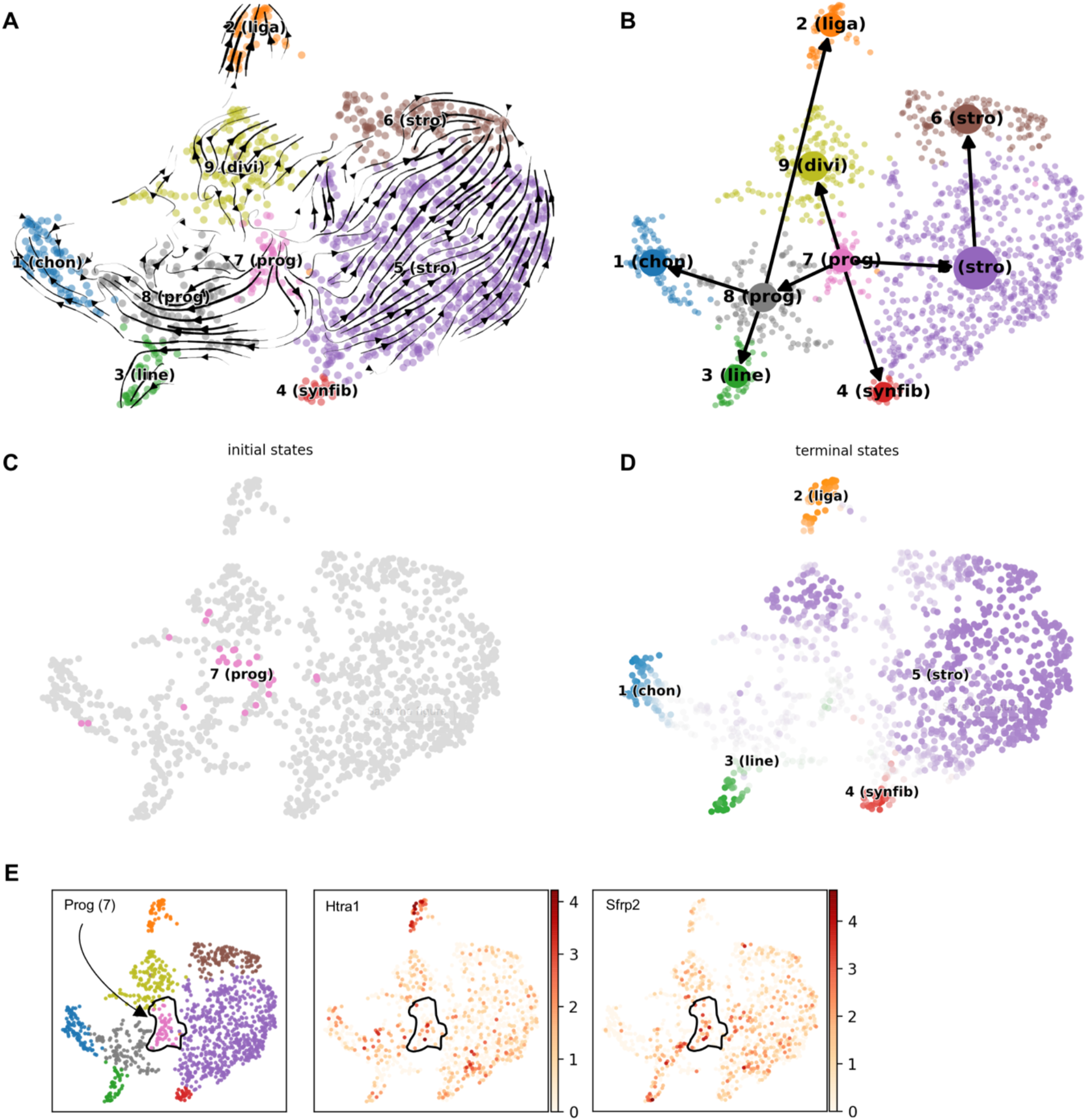
Developmental relationship among GLE cells. (A) RNA velocity analysis. Arrows indicate the predicated future state of cells. (B) PAGA analysis. Arrows summarize the RNA velocity results between clusters. CellRank identifies initial (C) and terminal (D) states of cell fate potential. Cells are colored by states. (E) Expression of interzone genes Htra1 and Sfrp2 in progenitor cluster 7.

To summarize these patterns of transcriptional velocity, we used PAGA, which consolidates the individual cell trajectories into connectivity’s between clusters ^36^. PAGA analysis predicted that cluster 7 flows into the stromal lineages (clusters 5 and 6), the synovial fibroblasts (cluster 4). Cluster 7 also flows into cluster 8, which subsequently flows into the chondrocyte, lining, and ligament clusters (**Fig 5B**). This suggested that the cells of clusters 7 and 8 continued to serve as a reservoir of progenitor cells. To explore this possibility more rigorously, we used the CellRank computational tool, which uses a combination of RNA velocities and cell-cell similarities to infer fate potential in scRNA-Seq data ^35^. CellRank identified cluster 7 as an initial cluster from which cells traverse trajectories towards terminal states represented by cluster 1 (chondrocytes), cluster 2 (ligamentocytes), cluster 3 (lining cells), cluster 4 (synovial fibroblasts), and cluster 5 (stromal cells) (**Fig 5C-D**). It is possible that cluster 7 progenitor cells are residual, or late differentiating, interzone cells as they do express detectable levels of genes Htra1 and Sfrp2 (**Fig 5E**) that are expressed at earlier time points in the interzone ^5^. However, this does not exclude the possibility that they are more recently immigrated Gdf5-lineage cells that primarily contribute to meniscus and intra-articular ligament ^37^.

### 3.4 FGF-MAPK signaling distinguishes early from late stages of GLE differentiation

Next, we sought to identify the signaling pathways that regulated transitions from the progenitor populations to each of the more differentiated end points: ligamentocytes, synovial fibroblasts, chondrocytes, and lining cells. To achieve this, we first performed differential gene expression analysis on the pairs of cell clusters that were predicted by RNA Velocity, PAGA, and CellRank to be immediately related to each other (**Fig 6A**). These pairs were 7 (prog) to 8 (prog), 7 (prog) to 4 (synfib), 8 (prog) to 3 (line), 8 (prog) to 2 (liga), and 8 (prog) to 1 (chon). Then we calculated the extent to which genes upregulated in each cluster (as compared to their immediate progenitor) were enriched in known targets of 18 effectors of nine signaling pathways crucial in development (FGF-MAPK, FGF-PI3K, FGF-STAT, Hedgehog, Hippo, Notch, TGFb-BMP, TGFb-Activin, and Wnt).

**Figure 6:**
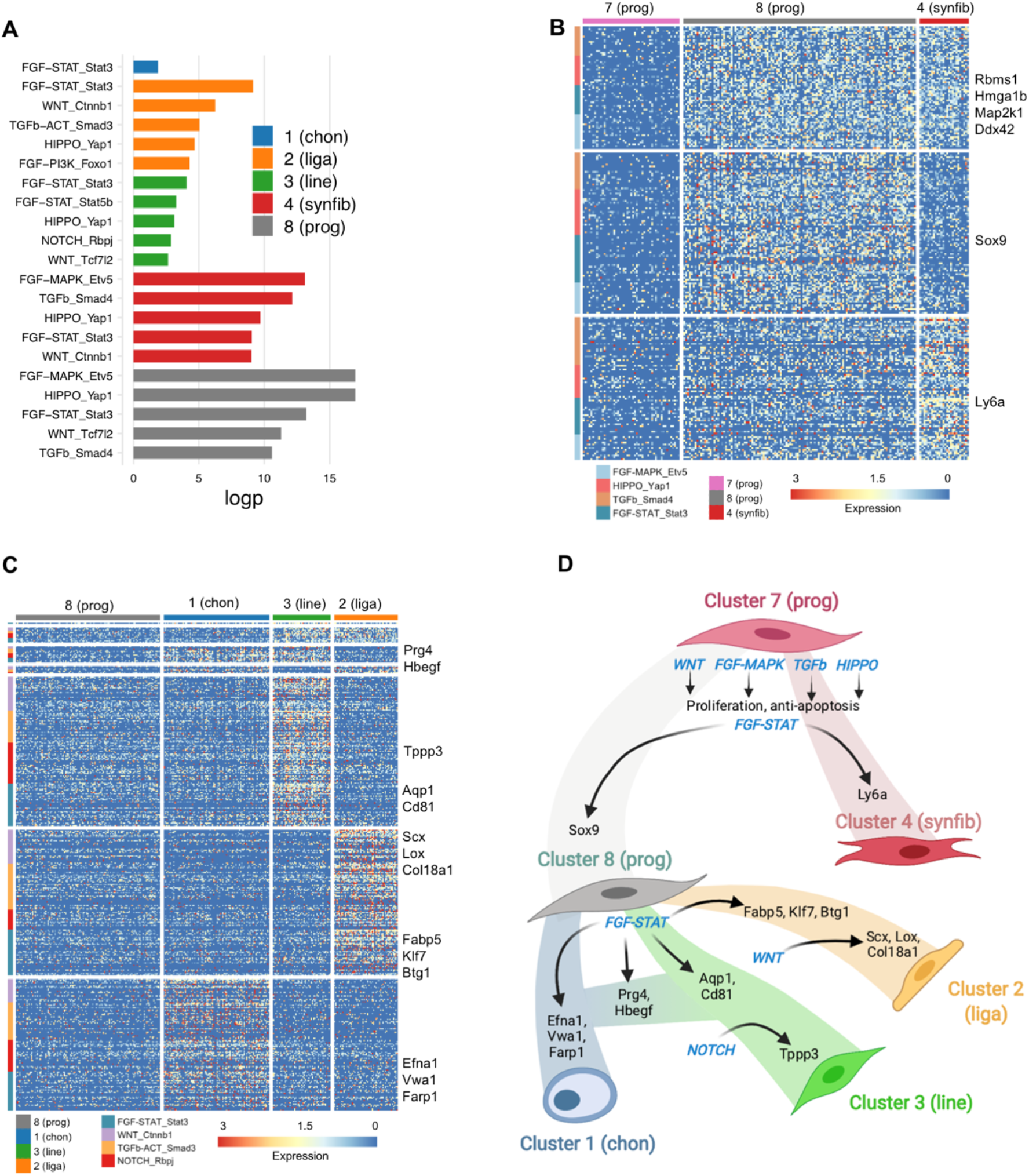
Signaling pathways that contribute to GLE cell differentiation. (A) Gene set enrichment of genes up-regulated in each cluster relative to the clusters predicted progenitor state. Gene signatures tested were gene sets of signaling pathway effector targets as determined by ChIP-Seq. Clusters 4 and 8 were compared to cluster 7. Clusters 1, 2, and 3 were compared to cluster 8. (B-C) Heatmap showing the genes of enriched signaling pathways in Cluster 8 vs 7 and Cluster 4 vs 7 (B), and Cluster 1, 2, or 3 vs cluster 8 (C). (D) Diagram of signaling pathways regulating transitions between indicated cell states.

We found that targets of Etv5, an effector of the FGF-MAPK pathway, and Yap1, the effector of the Hippo pathway, were highly enriched in both the progeny of cluster 7: cluster 4 (synfib) and cluster 8 (prog) (**Fig 6B**). Many of the enriched genes are associated with proliferation, such as Rbms1, Hmga1b, and Map2k1, consistent FGF-MAPK’s and Hippo’s role in controlling the size of progenitor pools ^38,39^. On the other hand, some signaling pathways were either only enriched in one progeny cluster (e.g. WNT_Ctnnb1 in the synovial fibroblast cluster 4) or were enriched in both but had distinct targets activated. For example, chondroprogenitor master regulator and FGF-STAT_Stat3 target Sox9 is only activated in the progenitor cluster 8, whereas the FGF-STAT_Stat3 target Ly6a is mainly activated in the synovial fibroblast cluster 4 (**Fig 6B**).

When we examined the pathways enriched in progeny of cluster 8, we again found pathways that were unique to each lineage, pathways that were enriched in more than one lineage, and pathways that were commonly enriched but had distinct target genes upregulated in different lineages. FGF-STAT_Stat3 targets were upregulated in chondrocytes, ligamentocytes, and lining cells compared to their progenitors in cluster 8. Both Prg4, which encodes lubricin, and the Egf ligand Hbegf, which is upregulated in osteoarthritis ^40^, are FGF-STAT_Stat3 targets upregulated in the lining and chondrocyte clusters (**Fig 6C**). FGF-STAT_Stat3 targets that are unique to each lineage include endothelial-associated Aqp1 and Cd81 in lining cells; Fabp5, Klf6, and Btg1 in ligamentocytes; and Efna1, Vwa, Farp1 in chondrocytes. Interestingly, the chondrocyte cluster was not marked by enrichment of any other signaling pathway, suggesting that in the absence of other signaling events, it is the default fate from the progenitor 8 state. We also note that nascent ligamentocytes were marked by enrichment of WNT signaling and that the effector targets included the tenocyte/ligamentocyte regulator Scx, as well as Lox, and Col18a1, which are genes encoding proteins important to the structural integrity of ligament ^41^. A summary of our analysis of signaling pathways and the targets of their effectors is depicted in **Figure 6D**.

### 3.5 Identification of transcriptional circuits underpinning joint population diversity

Many lineage specific genes were not predicted to be directly regulated by effectors of the signaling pathways that we analyzed. In addition to signaling pathways, cell intrinsic gene regulatory networks (GRN) contribute to cell fate choice and differentiation during development ^42^. To identify the GRNs that underpin joint cell diversification and differentiation, and in particular to identify the regulators of lineage specific genes, we used Epoch, which leverages pseudotemporal ordering to infer dynamic GRNs ^28^. Epoch defines lineage- or trajectory-specific GRNs, and it divides these temporally into time periods, or epochs, to identify dynamic regulatory relationships. The rationale behind this approach is that developmental GRNs change as cells differentiate such that transcription factors can regulate different genes in different lineages and at different stages of development within a lineage.

We used Epoch to reconstruct the GRNs governing the transitions from the progenitor cluster 8 to the chondrocyte cluster 1 and the lining cell cluster 3. We were particularly interested in identifying the biological pathways that characterized each time period, their regulators, and the regulators of genes specific to each lineage (e.g. Col22a1 in the lining cluster and Col9a3 in the chondrocyte cluster) and genes shared in both lineages (e.g. Hbefg and Prg4). Therefore, we first used Epoch to identify the major time periods that marked the progression from progenitor cluster 8 to the chondrocyte cluster 1, and we performed gene set enrichment analysis on genes preferentially expressed in each resulting time period **(Fig 7A**). The early stage of chondrocyte differentiation was characterized by the extracellular matrix organization, regulation of insulin-like growth factor receptor signaling pathway, and skeletal system development GO Biological Process categories. The presence of both ECM-production genes and of insulin-like growth factor pathway genes is consistent with the observation that IGF activation enhances the synthesis of cartilage matrix, and our analysis indicates that this is an early event in articular cartilage differentiation ^43^. While the intermediate, transition stage was not enriched in any category, the final stage was enriched in cholesterol biosynthetic process, regulation of chondrocyte differentiation, and negative regulation of cell-substrate-adhesion. The activation of cholesterol metabolism programs is consistent with prior studies that link RORalpha expression to chondrocyte differentiation ^44^. The negative regulation of cell-substrate adhesion may play a role in how mesenchymal progenitors cells acquire the stereotypic spherical morphology of chondrocytes. Indeed, over-expression of Meltf, one in this pathway that is up-regulated in the final stage of the articular chondrocyte trajectory, in ATDC5 cells promotes a more chondrocyte-like shape and promotes differentiation ^45^.

**Figure 7:**
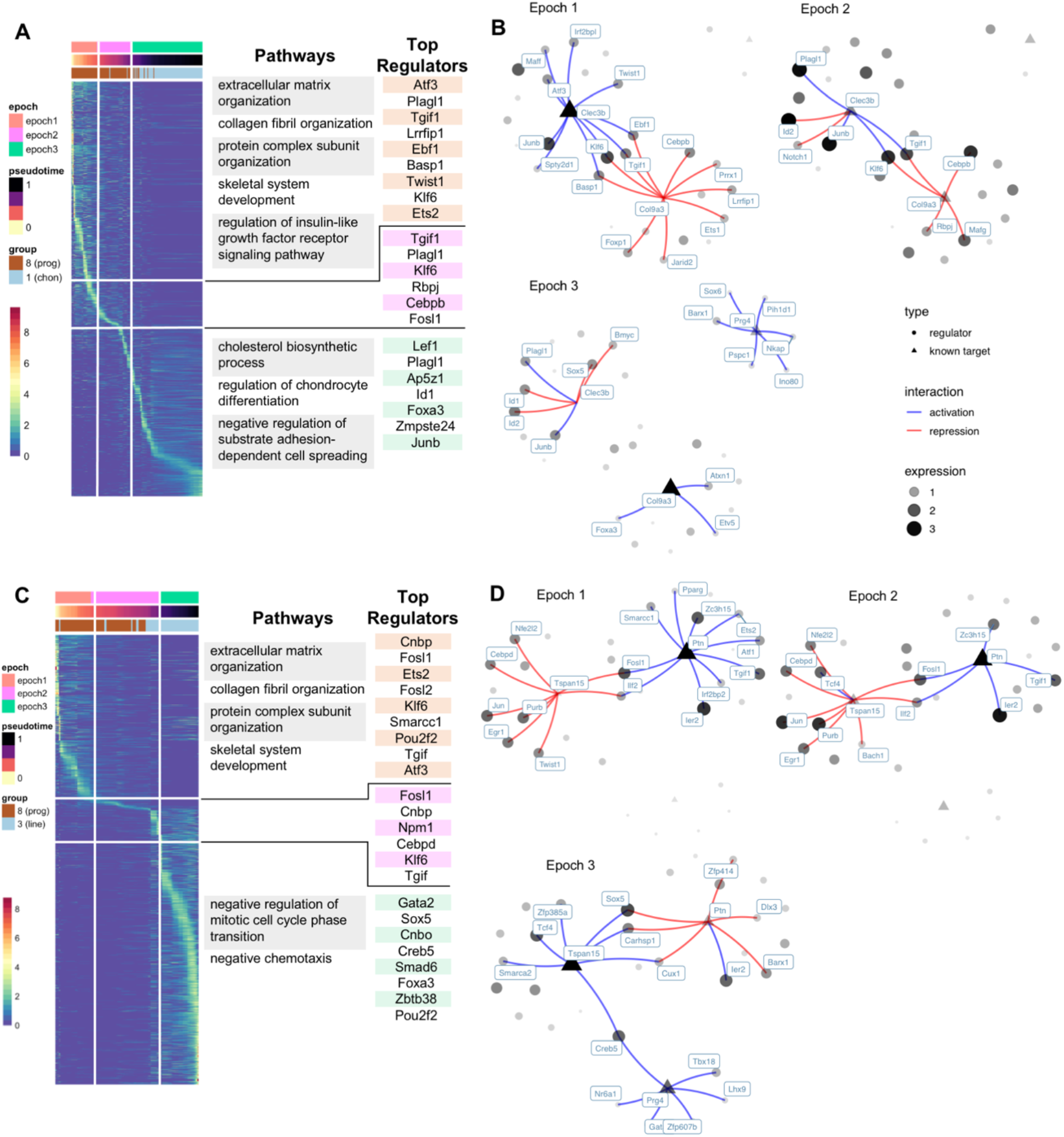
Dynamic GRNs that govern the transition from progenitor to chondrocyte and lining cell. (A) Heatmap of genes dynamically expressed along the chondrocyte trajectory. Epoch divides cells (columns) and genes (rows) into stages or epochs. Results of enrichment analysis of genes up-regulated in each epoch are shown to the right of the heatmap. The Epoch algorithm also reconstructs dynamic gene regulatory networks (GRNs). The top regulators of each epoch, determined by the network importance metrics of centrality and betweenness, are listed to the right of the enrichment results. (B) Sub-networks of that exemplify genes specific to the early (Clec3b) and late (Col9a3 and Prg4), and their regulators. (C-D) Similar to (A and B) but analysis performed on progenitor to lining cells trajectory.

Next, we sought to identify the regulatory circuits that contribute to the overall chondrocyte trajectory and to identify the transcription factors that directly regulate articular chondrocyte specific genes. Therefore, we used Epoch to infer the dynamic GRNs associated with the chondrocyte differentiation trajectory. To identify the most influential transcription factors at each stage, we computed betweenness and degree, which measure the centrality and number of direct neighbors that a transcription factor has in a GRN, respectively (**Supp Fig 4**). This analysis revealed identified Plagl1, which has previously been documented as being co-localized at sites of chondrogenesis ^46^, as a prominent regulator at all three stages, albeit as an activator of early stage genes such as Col3a1, Arid5b, and Sfrp2 but a repressor of late stage genes such as Timp1, Slc1a5, and Hbegf (**Supp Table 2**). Tgif1 followed a similar pattern, as it is predicted to promote expression of early stage genes such as Aspn, Clec3b, Osr1, Ptn, and Vim, but repress later stage genes such as Acan, Col11a1, Col2a1, and Col9a1 (**Supp Table 3**). The top overall regulator of the final stage was the Wnt effector Lef1, consistent with its documented role in specifically promoting a superficial articular chondrocyte phenotype ^47^.

Finally, we sought to better understand how genes indicative of the progenitor stage and the later chondrocyte stage were directly regulated. We chose to examine the regulators of Clec3b, Col9a3, and Prg4 as exemplars of the progenitor and articular chondrocyte stages (**Fig 7B**). In the early stage, expression of Clec3b, which encodes a heparin-bind protein associated with osteoarthritis ^48,49^, is promoted by Atf3 and the mesenchymal regulator Twist1, suggesting that the up-regulation of Clec3b in OA is a re-activation of a latent developmental program that contributes to collagen fibril synthesis. Several other TFs that promote Clec3b expression are also predicted to repress Col9a3, including Tgif1, Ebf1, and Klf6. The intermediate stage is characterized by loss of regulatory interactions, both the activating influence of TFs on Clec3b and factors that repress expression of Col9a3. In the final stage, Clec3b is repressed by several TFs including Sox5 and the transcriptional repressors Id1 and Id2. On the other hand, the factors that repress Col9a3 are lost by the late stage whereas TFs that promote its expression, Foxa3, Atxn1, and Etv5, are all active in the last stage. Our analysis did not predict any repressive factors for Prg4. Rather, its up-regulation seems to be controlled by the activation of a cohort of TFs in the late stage, including Sox6 and Barx1, as well as several genes that encode proteins involved in chromatin remodeling (e.g. Ino80, and Pih1d1) and nuclear paraspeckles (e.g. Pspc1), which would be consistent with a model of repression by nucleosome occlusion of Prg4 regulatory regions.

Next, we performed a similar series of analyses to the differentiation trajectory of the lining cells (cluster 3). The early stage was enriched in largely the same GO categories as in the chondrocyte trajectory (e.g. extracellular matrix organization, collagen fibril organization, and skeletal system development), the intermediate stage also lacked enriched GO categories, but the final stage was enriched in negative regulation of mitotic cell cycle phase transition and negative chemotaxis (**Fig 7C**). There were similarities and differences in the most influential regulators of the lining trajectory as compared to the articular chondrocyte trajectory, too. For example, as in the chondrocyte trajectory, Atf3 and Tgif were top regulators of the early stages of the lining trajectory. However, Cnbp, a transcription factor implicated in craniofacial development and predicted to promote proliferative programs ^50^, was only found to be a top regulator in the lining trajectory where it was predicted to up-regulate early and intermediate stage genes. The top regulators of the final stage of the lining trajectory included Gata2, which was previously implicated as a repressor of MSC fate commitment ^51^, Sox5, the loss of which ablates Prg4 expression in lining cells ^52^, and Creb5, which promotes Prg4 expression in the superficial zone ^53^. Finally, we sought to better understand how genes indicative of the progenitor stage and the later lining stages were directly regulated. We chose to examine the regulators of Ptn, Tspan15 and Prg4 as exemplars of these stages (**Fig 7D**). Expression of the early stage marker Ptn was promoted by several TFs common to the early stage of the chondrocyte trajectory, including Tgif1 and Ets2. During the intermediate stage, the lining marker Tspan15 was remained largely repressed by TFs such as Cebpd and Egr1, however, the Wnt effector Tcf4, which is predicted to promote Tspan15, became active. By the final stage, Ptn was repressed by a cohort of TFs including Barx1 and Sox5, Tspan15 expression was promoted by Sox5, Tcf4, and Creb5, and Prg4 expression was promoted by Creb5, Tbx18 and Gata2. The top predicted regulators of Prg4 in the lining cells (i.e. Creb5, Tbx18, and Gata2) differed from those in chondrocytes (i.e. Barx1, Sox6, and Pih1d1), consistent with the idea that the regulatory programs needed to activate transcription of the same target gene vary by epigenomic context.

## 4. Conclusions

In summary, we have identified nine groups of GLE cells by scRNA-seq, including chondrocytes, superficial lining cells, ligamentocytes, synovial fibroblasts, fibrochondro-progenitors, stromal cells, and dividing cells. Differentiation from the early progenitor (Cluster 7) stage involved activation of WNT, FGF-MAPK, TGFb, and HIPPO signaling pathways (**Fig 6D**), and targets of the Wnt effector Tcf7l2 were enriched in the fibrochondro-progenitors (cluster 8) whereas targets of Ctnnb1 were enriched in the synovial fibroblast cluster. Furthermore, signaling through the same pathway had distinct effects on these two lineages: the FGF-STAT cascade up-regulated Sox9 in cluster 8 cells but upregulated Ly6a in synovial fibroblasts. Many signaling pathways were detected as enriched in differentiation of the fibrochondro-progenitor cluster towards the ligamentocyte lineage and the lining cell lineage, whereas the articular chondrocyte lineage was enriched only in the FGF-STAT pathway suggesting that it is the default fate of these progenitors. Finally, dynamic GRN reconstruction identified Atf3, Plagl1, Tgif1 as major regulators of the chondrocyte differentiation trajectory, and Cnbp, Fosl1, and Gata2 as major regulators of the lining cell differentiation trajectory. In conclusion, our study will be a valuable resource for the community to further explore the gene signatures, signaling pathways, and regulatory networks associated with synovial joint development and how they relate to diseases such as osteoarthritis. We have made our data and analysis results available for the community to explore at https://e17-mouse.herokuapp.com

## Acknowledgements

We thank Wenyan Lu for technical support with in situ hybridization probe designs. 10× Genomics single cell processing and sequencing was conducted by Melissa Olson, Kakali Sarkar and David Mohr at the Genetic Resources Core Facility, Johns Hopkins Institute of Genetic Medicine. Flow cytometry and cell sorting was conducted by Hao Zhang at the Cell Sorting Core Facility, Bloomberg School of Public Health, Johns Hopkins University. The work is supported by National Institutes of Health R35GM124725; National Science Foundation Graduate Research Fellowship under Grant No. DGE-1746891; Maryland Stem Cell Research Fund 2017-MSCRFF-3910 (Award ID: 90074850)

---

**Supplemental Figure 1:**
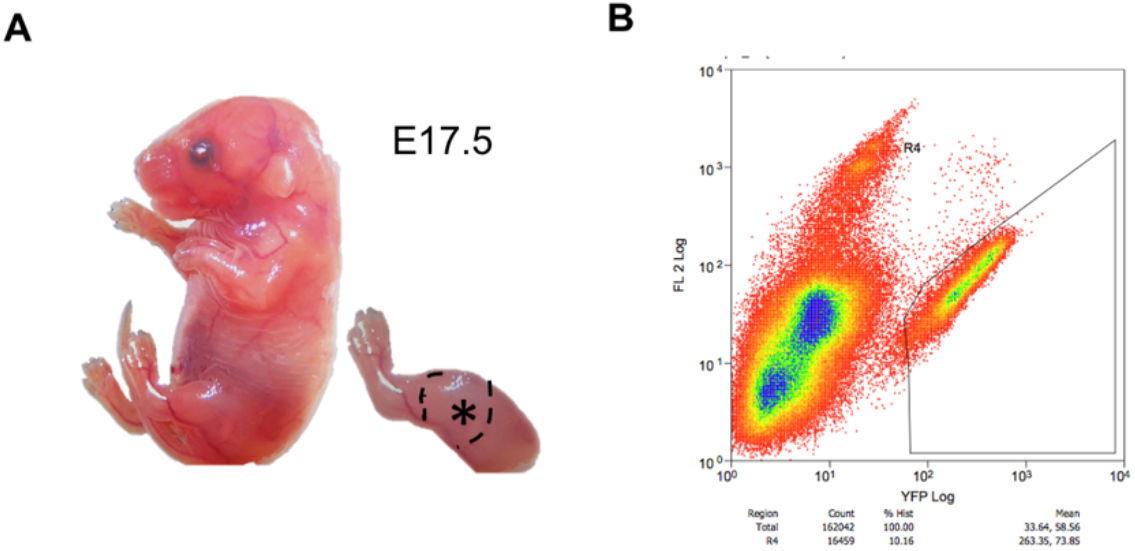
YFP^+^ cells collection. (A) E17.5 mouse embryo and star labels the region of hind limb dissected for cell isolation. (B) YFP^+^ cell isolation by FACs.

**Supplemental Figure 2:**
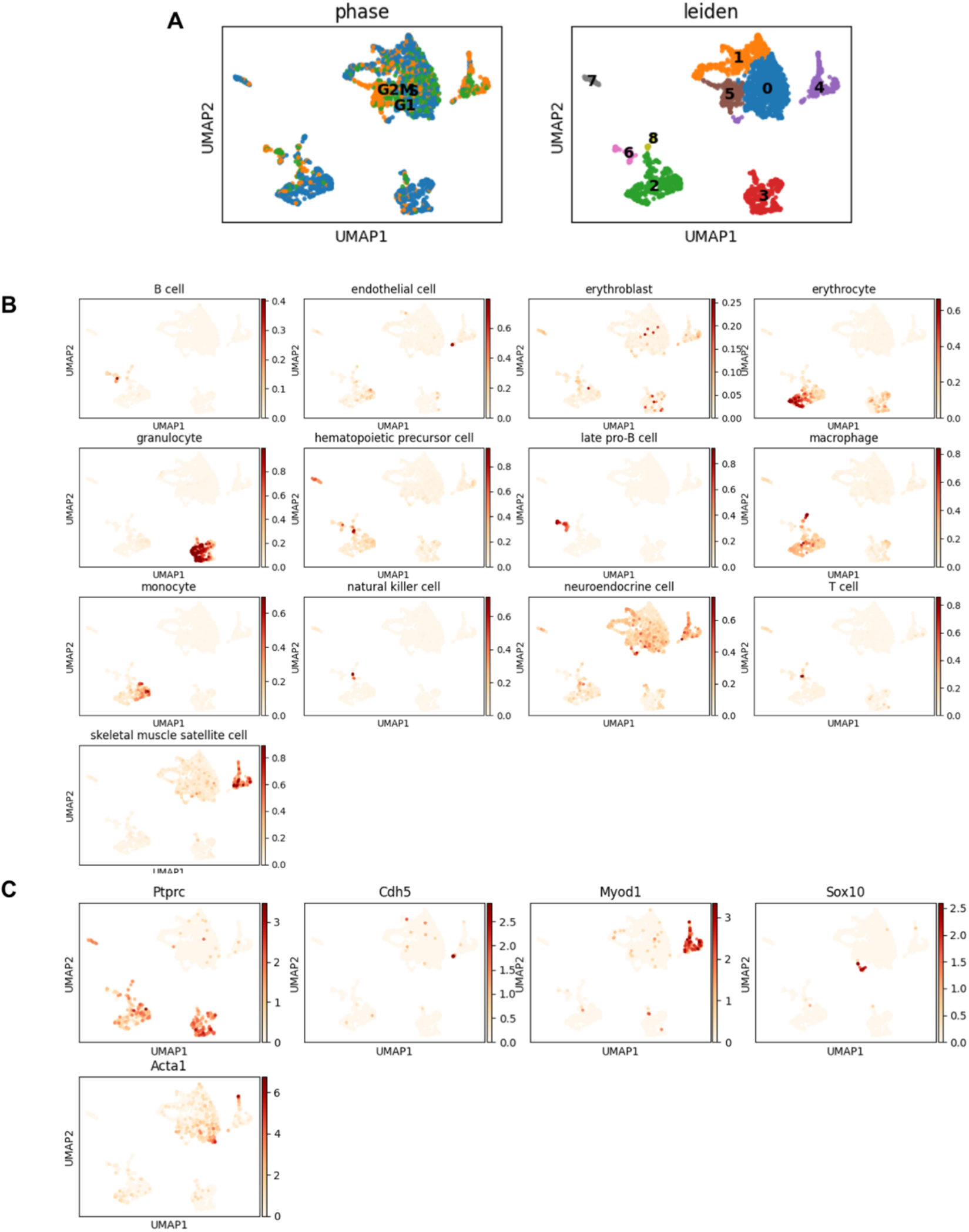
Initial clustering and identification of non-joint cells. (A) Leiden clustering and UMAP embedding of 9 groups of 2,468 cells, colored by mitosis phase (left), groups (right). (B) Cell types annotated by SingleCellNet. (C) Expression of five marker genes (Ptprc for blood cells; Cdh5 for endothelia cells; Myod1 for muscle cells; Sox10 for neural cells; Acta1 for smooth muscle cells).

**Supplemental Figure 3:**
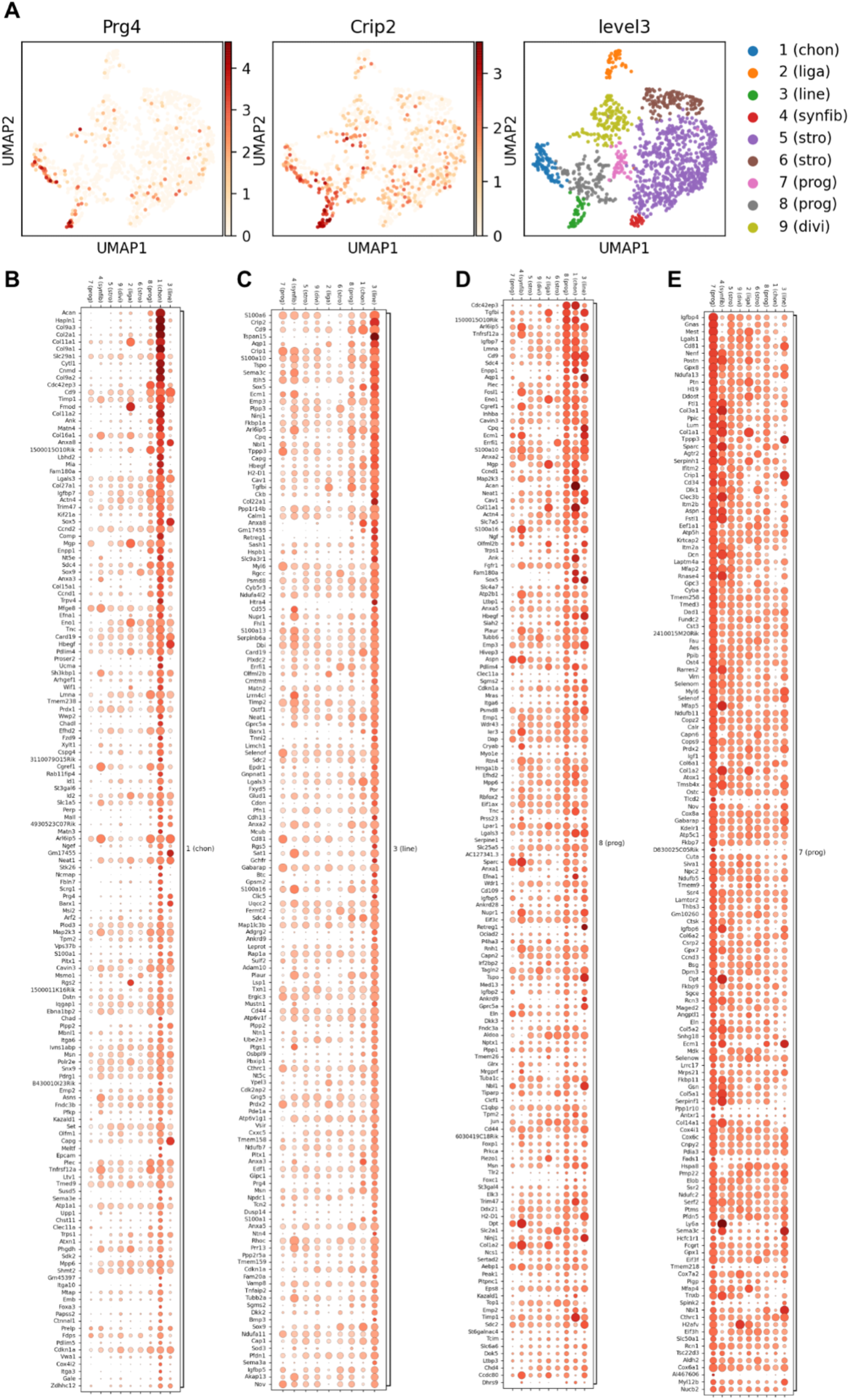
Identification of each cluster. (A) Gene expression pattern of Prg4 and Crip2. (B-E) Dot plots of 150 genes preferentially expressed in chondrocytes (B), lining cells (C), progenitors (cluster 8) (D), and progenitors (cluster 7) (E).

**Supplemental Figure 4:**
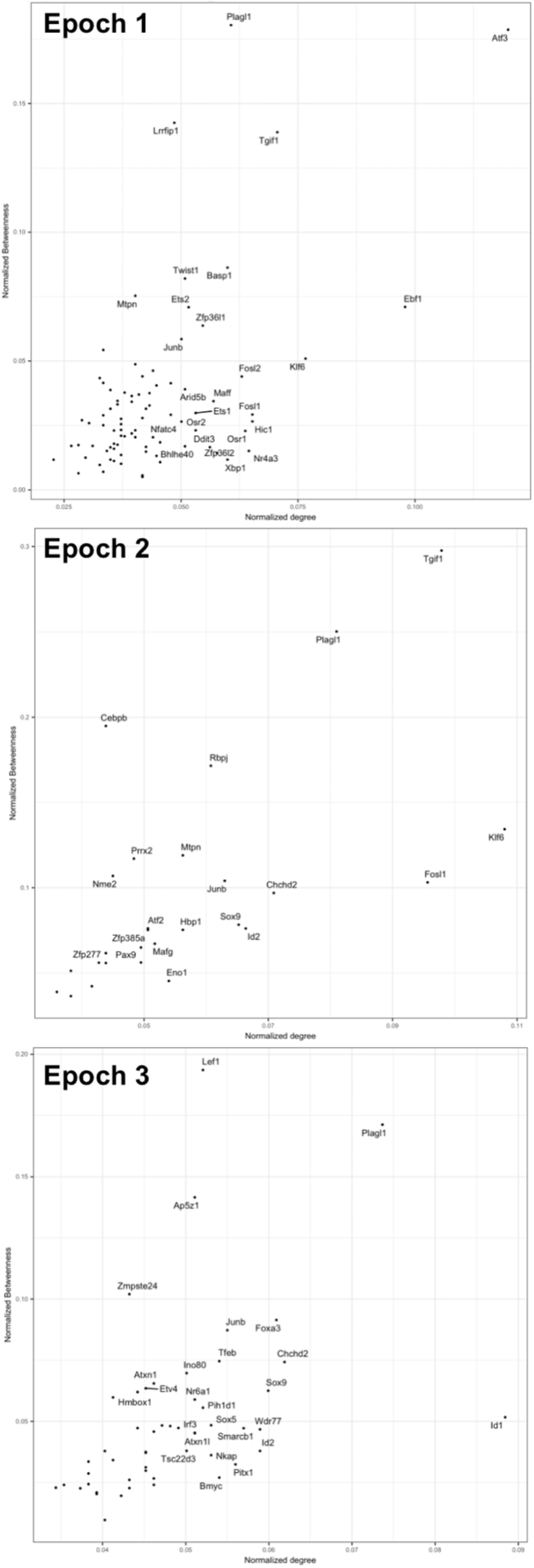
Regulators of Prog (cluster 8) to chondrocyte (cluster 1) trajectory.

**Supplemental Figure 5:**
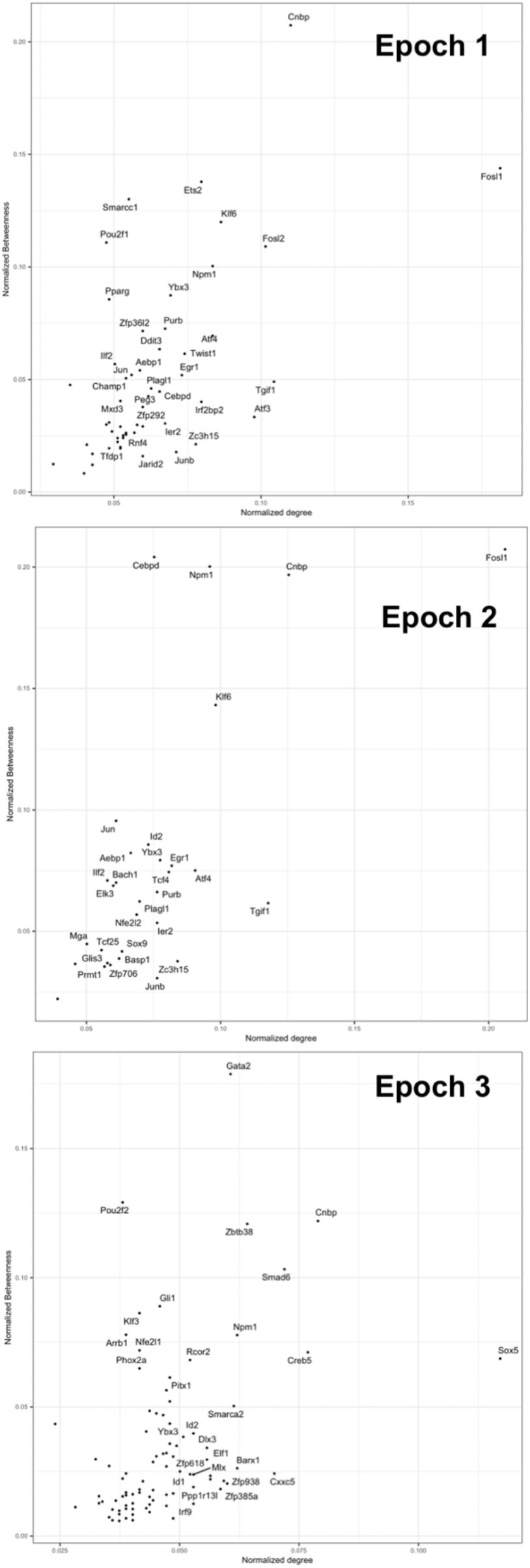
Regulators of Prog (cluster 8) to lining cell (cluster 3) trajectory.

**Supplemental Table 1:** Statistics on cells collected for scRNA-Seq. ‘Cells captured’ is determined by 10X CellRanger. GLE cells indicate the number of cells remaining after excluding cells unlikely to be GDF5-lineage, including hematopoietic cells, myoblasts, neural crest derived cells, endothelial cells and smooth muscle cells.

| No. YFP+ cells | % YFP+ cells | No. Captured cells | No. GLE cells | No. reads | Median reads per cells | Median Genes per cell |
| --- | --- | --- | --- | --- | --- | --- |
| 100.88K | 10.1% | 2648 | 1306 | 365,925,644 | 168,241 | 2,882 |

**Supplemental Table 2: GRNs of Prog (cluster 8) to chondrocyte (cluster 1)**. TG = target gene, TF = transcription factor, zscore = context-sensitive metric of association between TF and TG, corr = Pearson correlation coefficient of expression between TF and TF. Offset = the amount of pseudotime that the TF profile must be shifted in order to reach a maximal correlation with the TG.

**Supplemental Table 3: GRNs of Prog (cluster 8) to lining cell (cluster 3)**. TG = target gene, TF = transcription factor, zscore = context-sensitive metric of association between TF and TG, corr = Pearson correlation coefficient of expression between TF and TF. Offset = the amount of pseudotime that the TF profile must be shifted in order to reach a maximal correlation with the TG.

